# Realistic time-lags and litter dynamics alter predictions of plant–soil feedback across generations

**DOI:** 10.1101/2024.01.25.577053

**Authors:** Suzanne X. Ou, Gaurav S. Kandlikar, Magdalena L. Warren, Po-Ju Ke

## Abstract

Plant–soil feedback is a critical process in natural plant communities. However, it remains unclear whether greenhouse-measured microbial effects manifest in natural systems with temporally separated growing seasons as classic experiments often overlook seasonal time lags and litter dynamics.

We modified the classic two-phase experiment to study plant–soil feedback for three Californian annual plant species. Our response phase used soil inoculum obtained either immediately after plant conditioning, after a six-month dry period with the conditioning plant removed, or after a dry period with the litter of the conditioning plant. We characterized soil bacterial and fungal communities in different treatments and employed recent advancement in plant–soil feedback theory to predict plant coexistence.

Temporal delays and the presence of litter caused distinct responses in the fungal and bacterial communities, resulting in divergent microbial compositions at the end of the response phases. The delayed response treatments also affected microbially mediated stabilization, fitness differences, and invasion growth rates differently across species pairs, influencing predictions of plant coexistence.

Our study highlights that the interplay between seasonal delays and litter dynamics prevents the direct extrapolation of plant–soil feedback measurements across multiple seasons, emphasizing the necessity of considering natural history when predicting microbially mediated plant coexistence.

## Introduction

The interactions between plants and soil microbes have gained increasing recognition as a pivotal force in shaping plant communities (Bever *et al*., 2010, van der Putten *et al*., 2013). The effects of these interactions on plant community dynamics are most commonly studied under the plant–soil feedback (PSF) framework, which captures the effects of bidirectional interactions in which plants simultaneously alter and are affected by the soil microbial community (Bever *et al*., 1997). To implement this framework in empirical studies, PSF is often quantified through two-phase experiments that separate the feedback process into distinct “conditioning” and “response” phases (Bever *et al*., 1997, 2012). Plant performances during the response phase are measured to predict how soil microbes influence plant coexistence (Crawford *et al*., 2019, Yan *et al*., 2022). However, despite a vast body of literature showing that soil microbes can exert strong controls over plant species coexistence, connecting the predictions from such two-phase studies to the observed dynamics of plant communities in nature remains challenging (Forero *et al*., 2019, Beals *et al*., 2020, Beckman *et al*., 2022, Png *et al*., 2023). A promising approach for addressing this challenge is to adopt the classic two-phase design to better reflect the natural conditions under which PSF arises in the field (Gundale & Kardol, 2021).

Greenhouse experiments of plant–soil feedback typically simplify the temporal dynamics of feedback by conducting the conditioning and response phases sequentially, without any temporal separation between them. While this design likely captures the effects of microbial feedback among plants growing concurrently, whether these same effects manifest in communities characterized by temporally separated plant growing seasons or in communities where time-lags occur between soil conditioning and its subsequent recolonization is less clear. For example, Esch & Kobe (2021) found that in a temperate hardwood forest, *Prunus serotina* adults cultivate a soil community that suppresses the growth of conspecific seedlings, but this suppressive effect erodes within months of plant death. Thus, the long-term consequences of soil conditioning are unclear if there are time lags between adult death and subsequent arrival/growth of seedlings in the conditioned soils, which is especially likely in plant communities characterized by dispersal and/or seed limitation (Ehrlén & Eriksson, 2000). Similarly, in systems where plant dynamics are highly seasonal, the conditioning effects that build up during one growing season may not translate directly to affect plants in subsequent growing seasons if the soil community is reshaped during the intervening period (Barnard *et al*., 2013). Such dynamics are likely to be especially relevant in Mediterranean-type annual plant communities frequently used in PSF experiments (e.g., Bonanomi *et al*., 2012, Siefert *et al*., 2019, Kandlikar *et al*., 2021), where winter growing seasons are punctuated by dry summers of plant senescence (Elmendorf & Harrison, 2009). Furthermore, recent theoretical studies have demonstrated that the temporal dynamics of plant–soil feedbacks can substantially alter predictions of microbially mediated plant coexistence (Ke & Levine, 2021, Miller & Allesina, 2021). Thus, both empirical and theoretical evidence suggests that incorporating the natural temporal dynamics of plant communities into studies of plant–soil feedback might enable more robust predictions of how soil microbes shape plant coexistence in nature.

Another aspect of the soil conditioning process that is largely overlooked in two-phase plant– soil feedback experiments is that, in nature, the soil microbial community is shaped not only by the active conditioning effects of plants as they grow but also by the dead tissue (i.e., litter) that plants deposit onto the soil. Specifically, recent literature has shown that plant litter of different species can influence microbial communities by introducing phyllosphere microbes to the soil (Whitaker *et al*., 2017, Fanin *et al*., 2021, Minás *et al*., 2021) and by releasing chemicals and nutrients that affect soil microbial community assembly (Veen *et al*., 2021). These litter-induced changes in the microbial community can subsequently result in different plant–soil feedback on the responding plants (Veen *et al*., 2019, Aldorfová *et al*., 2022). For example, in systems with distinct phenology or seasonality, using soil collected at the end of the growing season rather than after decomposition would fail to capture the full impact of litter dynamics. Despite the role of litter dynamics in shaping soil communities in all systems, this process is largely overlooked in plant–soil feedback experiments, which typically remove all plant material at the conclusion of the conditioning phase. Incorporating the role of litter in plant–soil feedback is thus an important step for bridging the gap between classic experiments and natural conditions.

To better predict the long-term consequences of plant–soil feedback in natural systems, we also need theoretically robust metrics to extrapolate greenhouse experimental results. The original theory of plant–soil feedback popularized a pairwise feedback metric that quantifies how soil microbes drive frequency-dependent stabilization (e.g., via host-specific pathogens; Bever *et al*., 1997, Eppinga *et al*., 2018). Recent theoretical advances have integrated plant–soil microbe interactions with modern coexistence theory (Kandlikar *et al*., 2019, Ke & Wan, 2020), which utilizes invasion growth rates to predict species coexistence (i.e., quantifying whether each plant can establish in its competitor’s monoculture equilibrium from low density; Turelli, 1978, Chesson, 2000). Specifically, plant coexistence requires the stabilizing effects of microbes to overcome microbially mediated fitness differences, with the former capturing how microbes benefit both plants by driving negative frequency dependence while the latter capturing how microbes disproportionately impact one plant species over the other (Kandlikar *et al*., 2019, 2021, Yan *et al*., 2022). Evaluating coexistence outcomes on the basis of species’ invasion growth rates can also yield important insights for elucidating the underlying interactions in experimental data (Grainger *et al*., 2019, Ke & Wan, 2020, 2023). Examining the impact of experimental manipulation through these theoretical metrics enables a more nuanced understanding of the pathways through which plant–soil feedback influences plant coexistence.

Here, we conducted an experiment to address two questions about the role of soil microbes in shaping plant coexistence in annual grasslands: (1) How do seasonal time lags and plant litter decomposition interact with the conditioning process to alter the soil microbial community? (2) How do these changes to the soil community scale up to impact the predicted consequences of plant–soil feedback? To address these questions, we modified the two-phase greenhouse experiment and conducted three fully factorial response treatments. These treatments used soil inoculum obtained either immediately after plant conditioning, after a six-month dry period time lag with the removal of the conditioning plant, or after a similar dry period with the litter of the conditioning plant left intact. We quantified the absolute abundance of soil bacterial and fungal communities at the end of the conditioning and response phases, enabling us to evaluate how the soil inocula for each response treatment triggered different microbial communities. We then employed modern coexistence theory to predict the consequences of plant–soil feedback based on microbially mediated stabilization, fitness difference, and invasion growth rates (Kandlikar *et al*., 2019). Our results demonstrated that both time lag and plant litter altered the outcome of plant–soil feedback, with varying effects across species pairs. This work underscores the need to incorporate natural history when predicting microbially mediated plant coexistence.

## Methods

### Study system

We focused on three native Californian winter annual plants: a legume *Acmispon wrangelianus* (ACWR; Fabaceae), a grass *Festuca microstachys* (FEMI; Poaceae), and a forb *Plantago erecta* (PLER; Plantaginaceae). In spring 2019, we collected seeds from the University of California Sedgwick Reserve in Santa Barbara County, California, USA (34*^◦^*41’ N, 120*^◦^*02’ W), where all three species co-occur. In this Mediterranean-type climate, annual plants complete their life cycle and senesce in the hot, dry summer lasting about six months (May-October mean temperature = 18.9*^◦^*C, mean monthly precipitation = 4.57 mm; data from 2014–2023). The new generation germinates following rain in the cool, wet winters (November-April mean temperature = 12.3*^◦^*C, mean monthly precipitation = 54 mm). In September 2020, prior to winter rains, we collected field soil from Sedgwick Reserve to serve as microbial inoculum. To ensure that the field microbial community was not pre-conditioned by species in our experiment, we collected soil from four locations where there were no individuals of our focal species within a 1 m radius. The soils were kept at 4*^◦^* and transported to the lab within 12 hours, where equal amounts of soil from each location were sifted through a 2 mm sieve and homogenized. Prior to the experiment setup, we subsampled the field soil and stored it at -80*^◦^* for later DNA sequencing of the microbial community. One fraction of the field soil was then used to inoculate the conditioning phase pots, and the remainder was stored at 0*^◦^*C until further use in the response phases as a reference soil treatment.

### Greenhouse experiment and soil sampling

We modified the classic two-phase experiment to study how seasonal time lag and plant litter affect the soil microbial community and plant competitive outcomes. Specifically, our growth chamber experiment consisted of three fully factorial response treatments, using soil inocula that went through different handling treatments to represent these natural history factors (Fig. 1). We collected soil samples at different stages of the experiment and characterized the microbial community by high-throughput sequencing. Plant competitive outcomes were predicted by measuring plant biomass performance at the end of the experiment.

**Figure 1.**
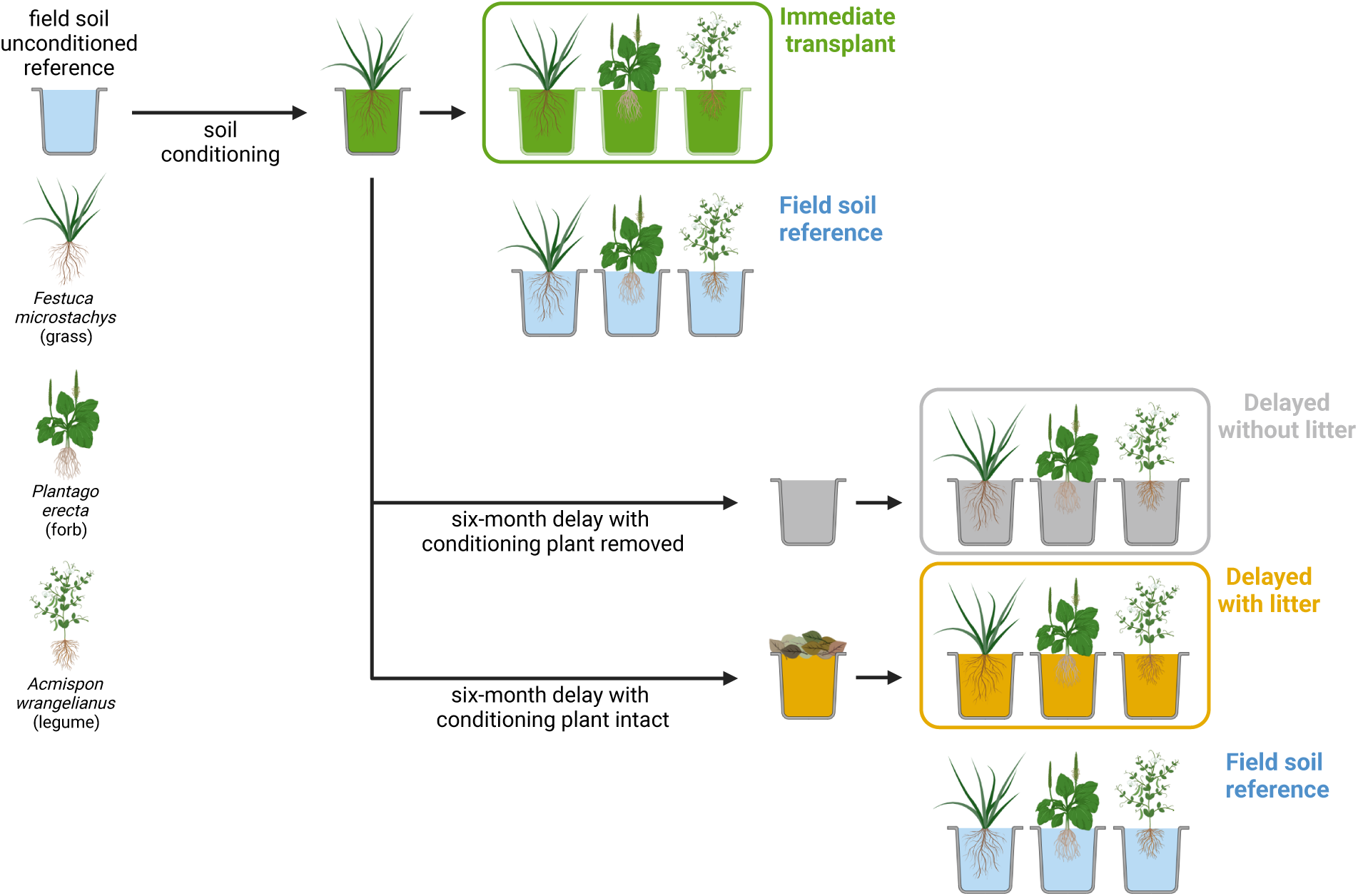
Schematic diagram of the two-phase plant–soil feedback experiment with three fully factorial response treatments: the immediate transplant treatment (light green pots), the delayed without litter treatment (grey pots), and the delayed with litter treatment (brown pots). The two rounds of response phases are six months apart, each including a treatment with field unconditioned soil as references (blue pots). Note that the sterilized soil treatments and batch control pots are not included in this illustration (see Methods).

#### Conditioning phase

To cultivate soil microbes associated with each species, we grew three high-density monocultures (8 g viable seed/m^2^) of each species in bleach-sterilized 1-gallon pots (Fig. 1). We first filled each pot with 2.60 L sterilized potting mix (equal parts sand, clay, peat, perlite, and vermiculite; autoclaved twice, each 2 hours with a 24-hour resting period in between). We then added 0.30 L of field soil to each pot and topped it with a 0.10 L layer of sterilized potting mix to achieve a 10% volume of live inoculum. Into each pot, we sowed 0.141 grams of seeds of a single species, which we had surface-sterilized by soaking in 1% bleach for 2 minutes and washing with ultrapure water twice for 1 minute each. We stored pots at 4*^◦^*C for five days to trigger germination, after which we moved pots to a growth chamber (25*^◦^*C, 60% humidity, 10:14 hour day:night cycle) for 80 days, approximately the length of a complete growing season. In addition to the 9 large conditioning pots, we grew 10 replicate individuals of each species in sterilized potting mix to serve as phytometers between the different phases of the experiment (3 species *×* 10 replicate individuals = 30 pots). We rotated control plants (30 pots) and conditioning monoculture pots (9 pots) weekly within the growth chamber.

The conditioning phase of the experiment concluded in December 2020. At this time, we randomly chose soil from one monoculture pot for each species to serve as the inoculum source for the “immediate” response treatment (green pots in Fig. 1). We designated the remaining two monoculture pots per species for the two time-lagged response treatments, and left these in the growth chamber (25*^◦^*C, 10% humidity, 10:14 hour day:night cycle) for an extra six-month dry period to mimic the temporal gap between two consecutive seasons. From one of these, we removed all aboveground biomass of the conditioning plant (grey pot in Fig. 1), whereas in the other we left all plant tissue intact (brown pot in Fig. 1). Thus, for each species, we were able to evaluate the effects of soil conditioning on subsequent plant growth without any time lag ("immediate” treatment), and could also evaluate how the presence of litter interacts with time lags to affect the plant performance during the “delayed” response phase (Fig. 1). Before using the conditioned monoculture pots for their corresponding response phase, we collected soil samples from each pot to characterize how seasonal time lag and plant litter influenced the soil microbial community (see section *DNA sequencing of the microbial community*).

#### Response phase

To create soil inocula for the “immediate” response phase, we removed the aboveground biomass from one conditioning monoculture pot per species, and sifted the soil through a 2 mm sieve to remove roots and homogenize the soil. We autoclaved half of this soil to create the sterilized inocula; the other half served as the live inoculum. We grew 10 replicate individuals of each species with either live or sterilized inoculum from each of the three species’ freshly conditioned monoculture pot (i.e., 3 plant species *×* 3 soil inoculum *×* 2 sterilization treatments *×* 10 replicates = 180 pots). These plants in immediate response treatment correspond to the typical experimental procedure in plant–soil feedback experiments, which use soil inoculum collected at the end of the conditioning phase. To quantify the impact of time lag and litter presence, we started another round of response phase (i.e., the two delayed treatments) in June 2021 using soil inoculum that experienced an additional six months of dry period after the conditioning phase, thereby more closely mimicking the natural temporal dynamics of these grasslands. For these treatments, we grew 10 replicates of each species with live and sterile inoculum from the corresponding “delayed” and “delayed with litter” monoculture pots of each of the three species (i.e., 3 plant species *×* 3 soil inoculum *×* 2 delay treatments *×* 2 sterilization treatments *×* 10 replicates = 360 individuals).

For the response phase pots, we filled 125 mL Deepots (Stuewe & Sons, Inc.) with sterilized potting mix and 10% volume of soil inoculum, and covered the inoculum with a thin layer of sterilized potting mix. We surface-sterilized and pre-germinated seeds, and transplanted seedlings into the pots so that each pot had a single individual. As evaluating microbially mediated coexistence outcomes requires growing plants in a reference soil (*sensu* Kandlikar *et al*., 2019), we also grew 10 replicate individuals of each species using unconditioned field soil as inoculum (previously collected and stored at 0*^◦^*C; blue pots in Fig. 1). To control for batch effects between the two rounds of response phases (i.e., the immediate response treatment in December 2020 and the two delayed response treatments in June 2021), during each round we grew 10 replicate individuals of each species with sterilized potting mix (i.e., batch effect controls with no inoculum). In total, the response phase of our experiment included 660 pots (i.e., 180 pots for the immediate response treatment + 360 pots for the two delayed response treatments + 3 plant species *×* 10 replicates *×* 2 rounds of field reference soil treatments + 3 plant species *×* 10 replicates *×* 2 rounds of batch effect controls). We grew the plants in a growth chamber for 80 days (same conditions as the conditioning phase, rotated weekly), after which we harvested, dried (72 hours at 60*^◦^*C), and weighed plant aboveground biomass. At the end of all response phases, we collected soil samples from each pot to characterize the soil microbial community.

### DNA sequencing of the microbial community

As described earlier, we collected soil samples from conditioned monoculture pots (before using them as soil inoculum) as well as response phase pots at the end of the experiment. The former was meant to characterize how seasonal time lag and plant litter affected the soil microbial community, while the latter was meant to see if these changes in the inoculum triggered long-lasting impacts (Hannula *et al*., 2021). We mixed soils within each pot well and subsampled 0.25 g of soil. To each sample, we added 8 ng of P and F synthetic chimeric DNA spike for the quantification of prokaryotic and fungal absolute abundance prior to DNA extraction (Tkacz *et al*., 2018). We extracted DNA using Qiagen DNeasy PowerSoil Pro Kit according to the manufacturer’s manual, with a 65*^◦^*C water bath incubation (10 minutes) prior to bead-beating to improve yield. Using twostep PCR, we amplified the V4 region of bacterial 16S ribosomal RNA gene (Caporaso *et al*., 2012) and fungal internal spacer 1 region (ITS1) (White *et al*., 1990) with index primers. We purified, normalized, and pooled amplicon libraries and sequenced using 2 *×* 300 bp paired-end Illumina MiSeq (see Supporting Methods S1).

### Data analysis

#### Microbial community

We converted raw binary base call (BCL) files to fastq files and demultiplexed with Illumina Bcl2fastq2 (v2.20). We trimmed adapter sequences from reads using cutadapt (Martin, 2011) with python (v3.10.9). Using the Divisive Amplicon Denoising Algorithm (dada2 v.1.28.0) in R (v4.3.0), we quality filtered and trimmed the fastq files, and inferred amplicon sequence variants (ASV) (Callahan *et al*., 2016a) following the published workflow (Callahan *et al*., 2016b). Specifically, we discarded low-quality ends of reads by trimming the bacterial forward reads to 250 bp and the reverse reads to 210 bp, discarding any reads shorter than these lengths. We chose not to trim fungal read lengths due to the varying size of the ITS gene region. We used decontam (Davis *et al*., 2018) to filter potential contaminant ASV and used phyloseq (McMurdie & Holmes, 2013) for downstream analysis. We discarded any ASV that was only detected in *≤* 5 samples; we also removed samples with extremely small or large read counts (i.e., more or less than 5x the average number of reads across all samples). We rarefied samples to 5000 sequencing reads for downstream analyses.

We transformed the community matrix from raw reads to relative abundance. We calculated absolute abundance as in Tkacz *et al*. (2018). Briefly, we divided the total number of environmental reads by the number of synthetic reads in each sample and then multiplied it by the number of gene copies in the 8 ng of synthetic spike-in, which was calculated by multiplying the number of gene copies in 1 ng of spike-in (i.e., 3.5E+07 for 16S and 1.2E+07 for ITS, respectively) by eight. To understand the starting soil microbial species pool that plants experience in the response phase, we explored differences in absolute abundances of 16S and ITS ASVs of the conditioned soil (immediately after the conditioning phase, after the six-month delay without litter, after the six-month delay with litter, and field reference soil). We first agglomerated the sequences in the conditioned soil samples to the class taxonomic level using tax_glom phyloseq function, and then visualized the total count for each class present in the samples. We identified compositional differences (i.e., beta diversity) with the Bray-Curtis dissimilarity metric and compared them with a permutational multivariate analysis of variance using the vegan (Oksanen *et al*., 2017) and stats R packages.

#### Plant biomass performance and competitive outcomes

To test for differences in plant biomass when grown with conspecific-conditioned versus reference soil (i.e., unconditioned field soil and sterilized potting mix) microbes, we conducted a series of linear models with log-transformed biomass values as the response variable, and soil source as the predictor. We fit separate models per species (ACWR, FEMI, and PLER) and treatment (immediate, delayed without litter, delayed with litter) to facilitate model interpretation. Prior to biomass analyses, we filtered out outliers within each experiment phase *×* species *×* soil *×* plant combination. Outliers were identified as individuals with biomass lower than Q1-1.5*IQR or higher than Q3+1.5*IQR, where Q1 and Q3 were the 25th or 75th quartiles, respectively, and IQR represents the difference between Q1 and Q3. We evaluated statistical significance at *α* = 0.05.

To predict how different response treatments modified the effects of plant–soil feedback on plant coexistence, we calculated microbially mediated stabilization and fitness differences for each treatment separately. We first quantified the effects of plant *j*-conditioned microbial community on the biomass performance of plant *i*, denoted as *m_ij_* (*i* and *j* = 1 or 2). This microbial effect is defined as *m_ij_* = ln(biomass of plant *i* in soil *j*) *−* ln(biomass of plant *i* in reference soil), i.e., the rate of plant *i* biomass accumulation when grown in soils with the plant *j* microbial community relative to that in an unconditioned reference soil. We then compare pots with conspecific versus heterospecific soil inoculum to calculate microbially mediated stabilization and fitness differences following the theoretical derivations in Kandlikar *et al*. (2019) (see also Kandlikar *et al*., 2021, Yan *et al*., 2022):

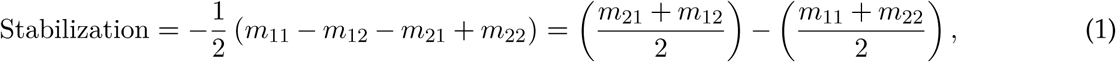

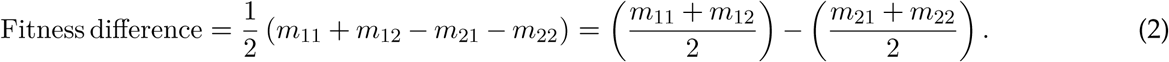

Here, microbially mediated stabilization (eqn. 1) quantifies how plants condition the soil to impact heterospecific relative to conspecific competitors. Positive values favor coexistence, as conditioned soils more negatively (or less positively) impact their hosts. Negative values (i.e., destabilization) indicate that conditioned soils more positively (or less negatively) impact host plants, and can drive priority effects. On the other hand, microbially mediated fitness difference quantifies how conditioned microbes disproportionately impact one plant species over the other: in the form of eqn. 2, a positive value indicates that plant 1 is favored by soil microbes because they benefit more from mutualistic microbes and/or suffer less from pathogenic microbes, and vice versa. We can thereby predict microbially mediated competitive outcomes based on these two metrics. If the absolute value of fitness difference overwhelms the absolute value of stabilization, then the plant with higher fitness will outcompete the other plant (i.e., plant 1 wins if eqn. 2 *>* 0 while plant 2 wins if eqn. 2 *<* 0). However, if the absolute value of stabilization exceeds that of fitness difference, then the theory predicts coexistence if eqn. 1 *>* 0 and priority effects if eqn. 1 *<* 0. Comparing eqns 1 and 2 across the three response treatments allows us to evaluate our hypothesis regarding how seasonal time lag and litter decomposition alter the coexistence consequences of plant–soil feedback.

As an alternative predictor of coexistence, we also calculated the invasion growth rate (IGR) of each species within a competing species pair. Specifically, the IGR of species 1 when growing in the monoculture equilibrium of plant 2 and its corresponding soil microbe, and vice versa, are as follows (Kandlikar *et al*., 2019):

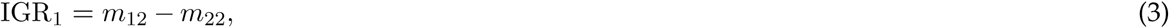

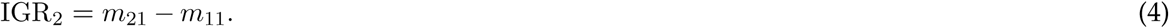

If the two IGRs differ in their signs, then the species with a positive IGR will outcompete the species with a negative IGR. If both IGRs are positive (i.e., eqns 3 and 4 *>* 0), the two species are predicted to coexist as both can recover from low density; alternatively, the theory predicts priority effects if both IGRs are negative (i.e., eqns 3 and 4 *<* 0). Furthermore, each species’ IGR only depends on how the resident soil microbe impacts the invader (e.g., *m*_12_), relative to their impact on the resident host (e.g., *m*_22_). Therefore, compared to eqns 1 and 2, which are aggregated metrics that incorporate the effects of both conditioned soil communities, it is easier to identify the key microbial impacts that are driving the changes in competitive outcome across response treatments.

We used a sampling approach to propagate the uncertainty when estimating *m_ij_* through to the predictions of plant competitive outcome (Yan *et al*., 2022, Terry & Armitage, 2023). Based on eqns 1–4, six different biomass terms are needed to predict the competitive outcome (i.e., the biomass of each plant species growing in soil 1, 2, and in the reference soil). For each sample draw, we randomly sampled one value for each of the six biomass terms from a normal distribution, with a mean equal to the empirical mean biomass and a standard deviation equal to the empirical standard error (SE). We repeated this procedure 1000 times for each species pair and calculated the stabilization, fitness difference, and invasion growth rates. This approach, compared to the commonly used orthogonal error bars (e.g., Kandlikar *et al*., 2021), better propagates uncertainty as it captures the interdependence between parameter estimations (Terry & Armitage, 2023). All analyses were conducted in R (v4.3.0) (R Core Team, 2021).

## Results

### Microbial community

We first present results of the microbial community in the soil inocula, which revealed that seasonal time lag and plant litter decomposition had a clear impact on bacterial and fungal abundance (Fig. 2). The bacterial abundance was highest in soils collected immediately following the conditioning phase (i.e., inoculum used for the immediate response treatment), and decreased in soils from each of the three plant species after a six-month delay period (i.e., inoculum used for the two delayed treatments; Fig. 2A). Fungal communities, which were much lower in absolute abundance, exhibited more variable patterns: total abundance increased after the six-month delay for soils conditioned by *P. erecta*, but declined for soils conditioned by *A. wrangelianus* (Fig. 2B). For soils conditioned by *F. microstachys*, the community pattern remains unclear as no fungal reads (aside from the synthetic spike-in) were detected in the delayed treatment without litter. Moreover, despite a notable proportion of unidentified taxa, we observed clear compositional shifts in the fungal community. For instance, Dothideomycetes were abundant in field reference soils but nearly absent in plant-conditioned soils. These results suggest that the responding seedlings were exposed to different microbial communities across the three response treatments, reflecting differences in the microbial species pool in the soil inocula.

**Figure 2.**
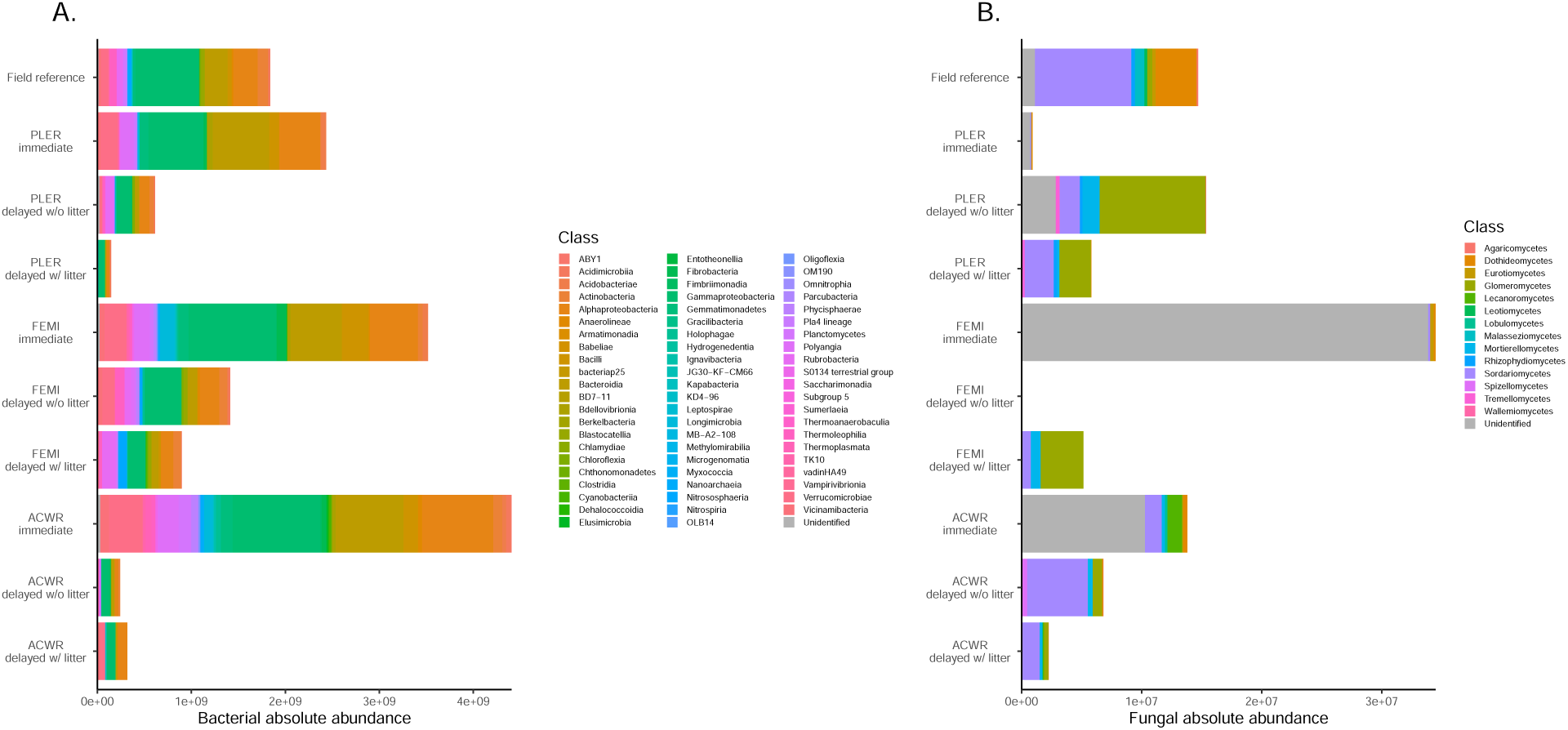
Absolute abundance of (A) bacterial and (B) fungal taxa (aggregated at the Class level) within the soil inocula used for different response treatments. Each stacked bar represents the absolute abundance of microbial taxa (x-axis and colors) within a specific soil inoculum (y-axis). The inocula are differentiated by their conditioning host species (i.e., *Acmispon wrangelianus* (ACWR); *Festuca microstachys* (FEMI); *Plantago erecta* (PLER)) and response treatments (i.e., immediate response, delayed without litter, and delayed with litter). The top row, labeled as field reference, depicts soil samples collected from Sedgwick Reserve at the experiment’s outset (i.e., prior to the growth of any conditioning individual). OTUs that were unable to be identified to the Class level are plotted in grey and labeled ”Unidentified”

To evaluate whether different soil inocula led to divergent microbial composition, we also sequenced the microbial community at the end of each response phase (Fig. 3). As absolute abundances were quantified, we combined bacterial and fungal communities when analyzing differences in microbial community composition. For all soil sources conditioned by different plant species, microbial composition varied between response treatments (*A. wrangelianus*: *R*^2^ = 0.205; *F. microstachys*: *R*^2^ = 0.204; *P. erecta*: *R*^2^ = 0.201; Reference soil: *R*^2^ = 0.136; all *P <* 0.05). The response treatment was remained a significant predictor for all soil inocula when the bacterial and fungal communities were analyzed separately (Fig. S1–S2).

**Figure 3.**
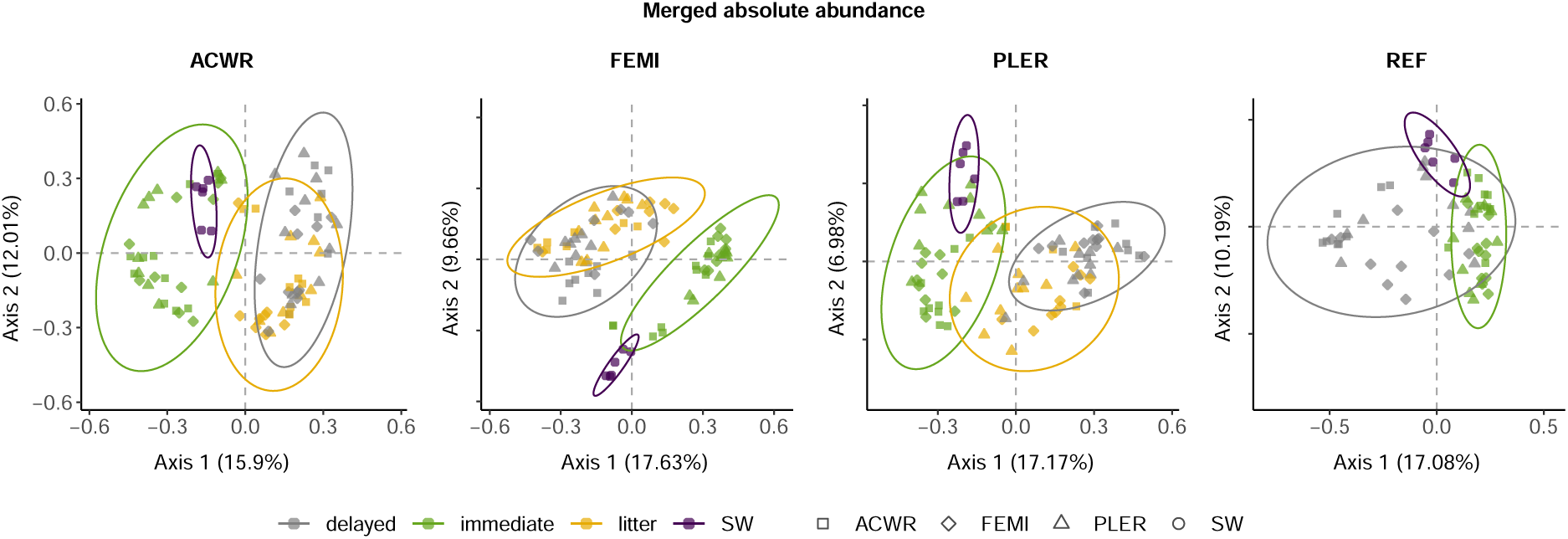
Principal coordinates analysis (PCoA) for the combined soil microbial community composition (i.e., bacterial 16S and fungal ITS) sequenced at the end of the response phase. Each panel represents a different inoculum source (conditioning host plant). From left to right: *Acmispon wrangelianus* (ACWR); *Festuca microstachys* (FEMI); *Plantago erecta* (PLER); unconditioned Sedgwick Reserve field soil as reference soil (REF). Each point represents the microbial community sampled from a seedling at the end of the response phase and the shape represents its species identity. Colors represent the three response treatments: immediate (light green), delayed without litter (grey), and delayed with litter (brown). As the two delayed treatments shared the same reference soil controls, we omitted one of the delayed treatment in the rightmost panel. Purple circles (labeled as SW) represent soils collected from Sedgwick Reserve at the beginning of the experiment (i.e., without the growth of any conditioning or responding individual) and were added for visualization purposes.

### Plant biomass

In the immediate response phase treatment, each of our focal species had lower aboveground biomass when grown with conspecific-conditioned soil microbes, relative to their growth in unconditioned field microbial community (ACWR: *F*_1,17_ = 4.89, *P* = 0.041; FEMI: *F*_1,16_ = 10.64, *P* = 0.049; PLER: *F*_1,17_ = 30.68, *P <* 0.001; Fig. 4; see also Fig. S3 for all biomass results). *F. microstachys* and *P. erecta* grew worse in soils with a conspecific soil community than in sterilized potting mix (FEMI: *F*_1,16_ = 7.21, *P* = 0.016; PLER: *F*_1,16_ = 9.53, *P* = 0.007), while the opposite was true for *A. wrangelianus* (*F*_1,16_ = 4.53, *P* = 0.048). Plants generally grew poorer in both delayed treatments (i.e., with/without litter present in conditioned soils during the time lag) compared to the immediate treatment (Fig. 4). Specifically, in the delayed treatments, *A. wrangelianus* grew substantially better when inoculated with any live soil microbial community than with sterilized potting mix, but its growth in conspecific-conditioned microbes was not significantly different than with an unconditioned field community (Fig. 4A). In contrast, *F. microstachys* growth in the delayed treatments was substantially lower with any live soil community than in sterilized potting mix. When litter was removed at the end of plant conditioning, conspecific-conditioned soil microbes resulted in lower plant biomass relative to unconditioned field microbes in *F. microstachys* and *P. erecta* (FEMI: *F*_1,14_ = 6.47, *P* = 0.02; PLER: *F*_1,18_ = 27.57, *P <* 0.001). This effect was diminished when plant litter was left intact after the soil conditioning phase.

**Figure 4.**
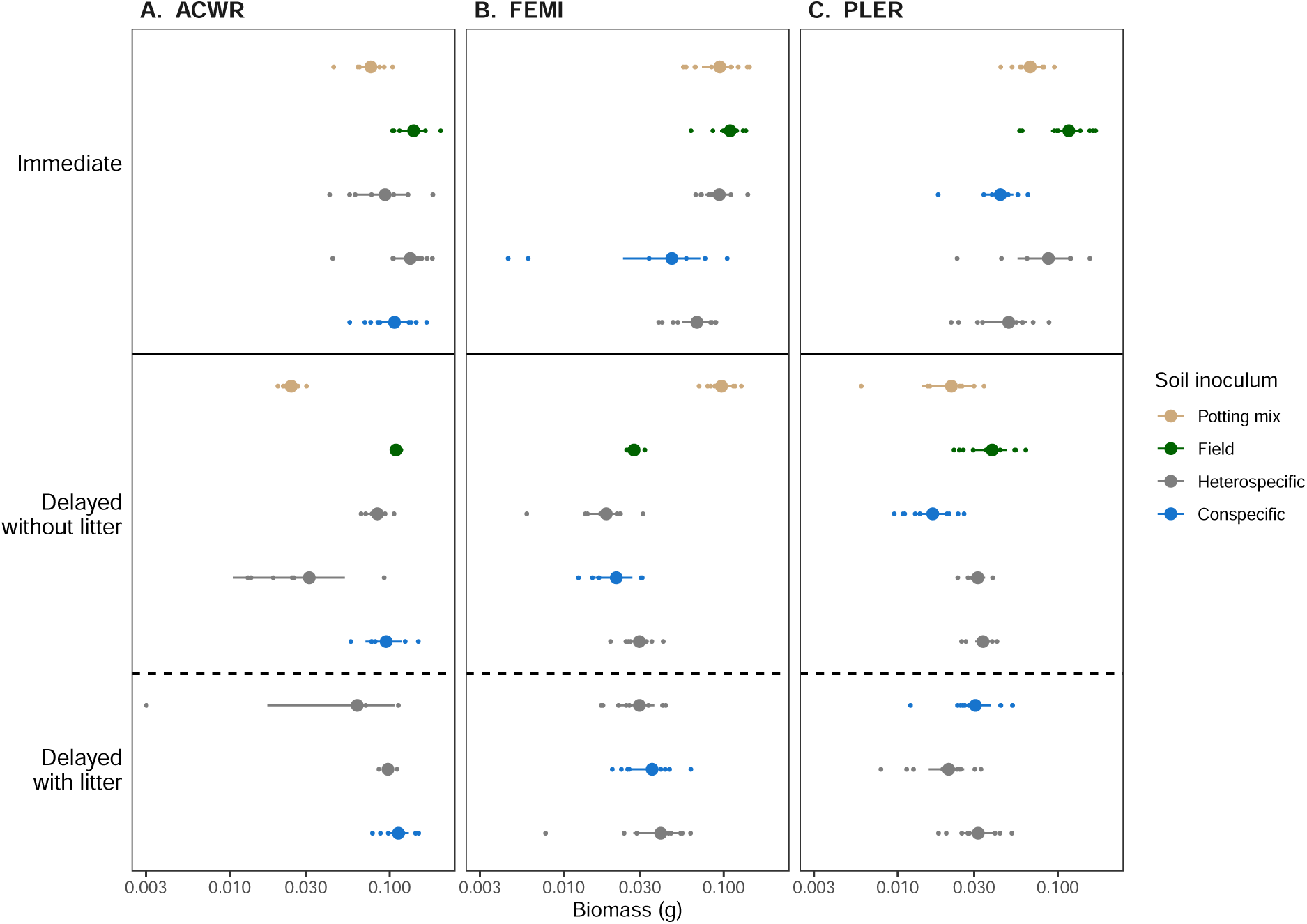
Effects of soil microbial inocula on plant biomass for (A) *Acmispon wrangelianus* (ACWR), (B) *Festuca microstachys* (FEMI), and (C) *Plantago erecta* (PLER). Each panel shows the aboveground biomass (log-scale x-axis) of the focal plant, grown with a soil microbial community that had been conditioned by conspecifics (blue) or heterospecifics (grey circles); unconditioned communities from field soil (green); or sterilized potting mix (brown). Note that the two delayed treatments shared the same field reference and sterilized potting mix controls. The three plant-conditioned soil inocula are ordered (from bottom to top) as follows: ACWR, FEMI, and PLER. Larger symbols indicate the mean biomass, error bars show 2 *×* SEM, and small points show each individual biomass.

### Plant coexistence outcomes

The aforementioned biomass differences across the three response treatments resulted in shifts in competitive outcomes for all three species pairs (Fig. 5). Although each species pair responded differently, the two delayed treatments mostly decreased the strength of stabilization (i.e., destabilization), with the exception being the delayed without litter treatment for the *A. wrangelianus*–*P. erecta* pair (Fig. 5C). The legume plant *A. wrangelianus* was predicted to outcompete *F. microstachys* and *P. erecta* in the immediate response treatment (green points in Fig. 5B–C). However, fitness differences shifted toward a direction that disfavor *A. wrangelianus* in the two delayed treatments (grey and brown points in Fig. 5B–C). Correspondingly, the IGR of *A. wrangelianus* became negative (Fig. 5E–F). As a result, *A. wrangelianus* loses its competitive dominance, exhibiting priority effect with *P. erecta* and being outcompeted by *F. microstachys* in the delayed with litter treatment.

**Figure 5.**
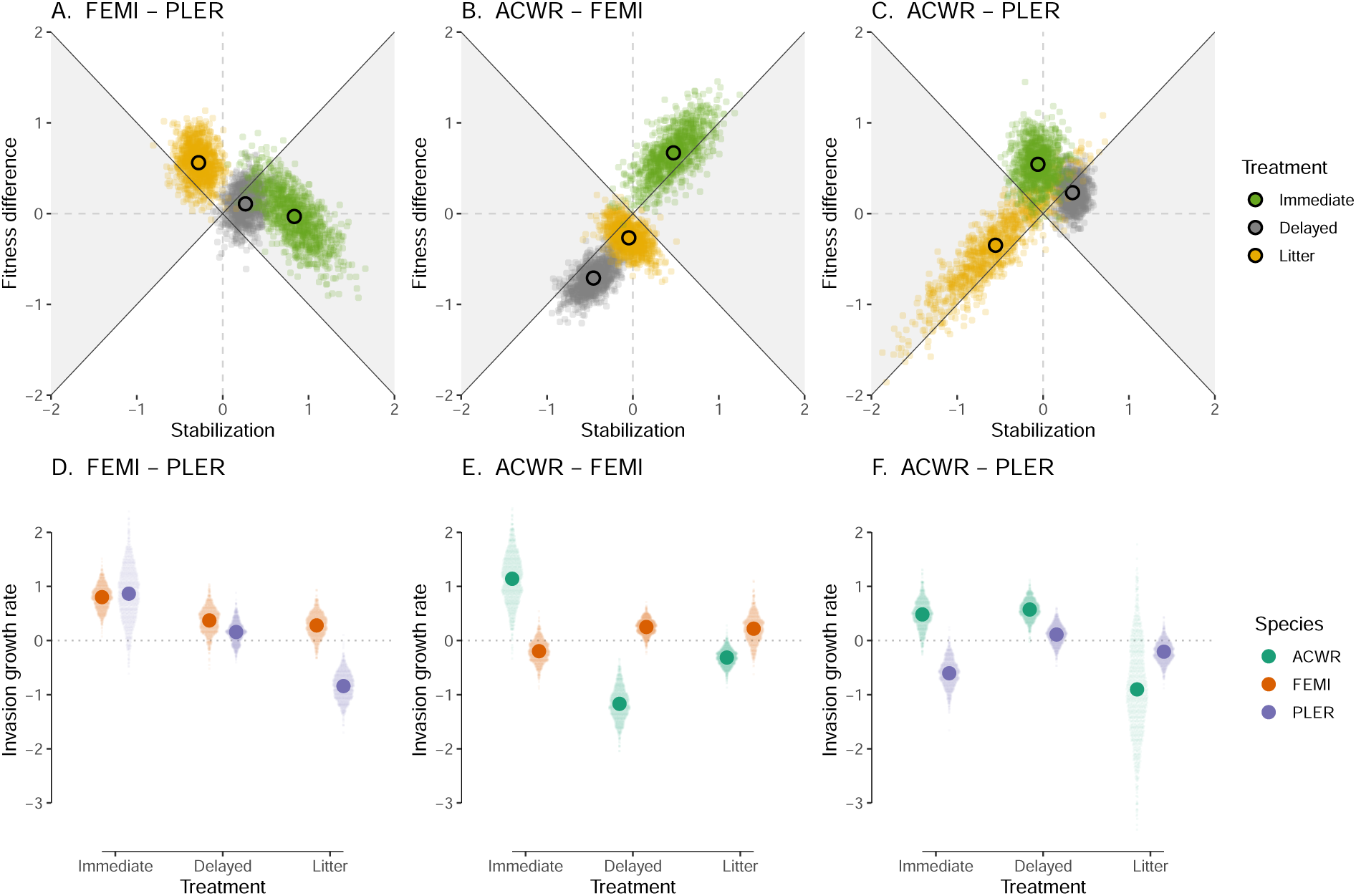
Predicted competitive outcomes between pairs of plants: (A & D) *Festuca microstachys* (FEMI) and *Plantago erecta* (PLER); (B & E) *Acmispon wrangelianus* (ACWR) and *F. microstachys*; (C & F) *A. wrangelianus* and *P. erecta*. For each panel, the first and second species listed on the facet label correspond to species 1 and 2 in eqns 1–4, respectively. (A–C) The parameter space of stabilization (x-axis) and fitness difference (y-axis) for the three species pairs. Each region represents different predicted competitive outcomes: the right and left grey triangular regions represent coexistence and priority effect, respectively. The upper and lower white triangular regions represent the dominance of species 1 and 2, respectively. For each species pair, the three response treatments are plotted on the same panel and are indicated by different colors: immediate (light green), delayed without litter (grey), and delayed with litter (brown). Each translucent point represents a random draw (see Methods) and the open black circle represents the mean stabilization and fitness difference of 1000 random draws. (D–E) Invasion growth rates (IGR, y-axis) for the three species pair under different response treatments (x-axis). Different colors represent different plant species: ACWR (green), FEML (orange), and PLER (purple).

We can also examine each species pair in more detail. For *F. microstachys* and *P. erecta*, the most common competitive outcome shifted from coexistence in the immediate treatment to *F. microstachys* outcompeting *P. erecta* in the delayed with litter treatment (Fig. 5A). In addition to destabilization, this shift in competitive outcome resulted from an increase in fitness difference in favor of *F. microstachys*. The corresponding decrease in the IGR of *P. erecta* (Fig. 5D) indicates that the shift in competitive outcome was mainly driven by changes in the soil microbes conditioned by *F. microstachys*. For *A. wrangelianus* and *F. microstachys*, the most common competitive outcome shifted from *A. wrangelianus* dominance in the immediate treatment to *F. microstachys* dominance in the two delayed treatments (Fig. 5B). This change in competitive outcome mostly resulted from the flip in the competitive hierarchy between the two plants (i.e., a decrease in fitness difference in favor of *F. microstachys*). A corresponding flip in the sign of the two species’ IGR can also be seen in Fig. 5E. For *A. wrangelianus* and *P. erecta*, the dominance of *A. wrangelianus* is the most common competitive outcome in the immediate treatment (Fig. 5C and E). The pair shifted towards coexistence (i.e., both plants have positive IGR) in the delayed treatment, but destabilization strengthened and resulted in priority effect (i.e., both plants have negative IGR) in the delayed with litter treatment.

## Discussion

Typical two-phase plant–soil feedback greenhouse experiments grow the responding plant immediately after soil conditioning in the greenhouse (Brinkman *et al*., 2010). When transplanted immediately, the predicted effects of soil microbes on species coexistence in our experiment were consistent with those of a previous study using the same system — the legume *A. wrangelianus* benefited from microbially mediated fitness advantage and was predicted to outcompete the other two species (Kandlikar *et al*., 2021) (light green points in Fig. 5). Yet during naturally occurring time lags of discrete growing seasons, the microbial community originally conditioned by plants may shift due to stochastic drift or biotic interactions between microbes. Moreover, the presence of litter can introduce microbes from other plant parts into the soil (Whitaker *et al*., 2017, Fanin *et al*., 2021) and the decomposition of litter can change the soil abiotic environment, thereby altering the soil microbial community (Veen *et al*., 2021, Minás *et al*., 2021). Thus, the microbial community encountered by a responding plant after the time delay may no longer resemble that of the original conditioning (Fig. 2). By quantifying stabilization and fitness differences, we show that microbially mediated plant coexistence outcomes change with the presence of a temporal delay and litter decomposition (grey and brown points in Fig. 5). Our study is a reminder that natural history, particularly in the form of temporal lags between consecutive generations, should be considered when designing and inferring long-term coexistence from plant–soil feedback experiments.

Our sequencing results showed a consistent decrease in bacterial abundance but a species-specific change in fungal abundance when comparing soils sampled immediately after conditioning to those sampled after the delayed treatments (Fig.2). Bacterial abundance was higher in the immediate response treatment relative to the field reference soil, indicating the proliferation of bacteria under plant conditioning in the growth chamber setting (Fig.2A). On the other hand, bacterial abundance decreased after the six-month delay, which may reflect drought-induced physiological stress, the lack of dormancy capacity, and intensified resource competition with other saprotrophic microbes (Shade *et al*., 2012, Schimel, 2018, Lennon *et al*., 2021). For fungal abundance, while communities conditioned by *A. wrangelianus* showed a similar decreasing pattern after the six-month delay, communities conditioned by *P. erecta* showed an opposite increasing pattern. We speculate that the initially low fungal abundance could be attributed to their slower growth rates, while the observed enrichment in the delayed treatments results from their greater ability to persist after host death (Challacombe *et al*., 2019). When comparing microbial community composition at the end of the response phases, we found that the total fungal and bacterial community diverged depending on response treatments (Fig. 3). This suggests that the temporal delay of previously conditioned soil has a long-lasting impact on the microbial species pool and their corresponding impact on the next generation.

Our results are the first to demonstrate the influence of a temporal delay by leveraging recent advancements in plant–soil feedback theory (Kandlikar *et al*., 2019). Moreover, our results highlight nuances when adopting modern coexistence theory to interpret empirical results. First, we show that quantifying invasion growth rates, in addition to the more commonly used stabilization and fitness difference metrics, yields a more nuanced understanding of the mechanisms through which microbial effects on plant coexistence arise. One example is the competition between *F. microstachys* and *P. erecta*, where the shift in competitive outcomes in the delayed treatments is caused by a loss of stabilization and an increase in fitness difference favoring *F. microstachys* (Fig. 5A). Examining the invasion growth rates suggests that the shift was primarily due to a decrease in the invasion growth rate of *P. erecta* (Fig. 5D), signifying a change in the microbial effects imposed by the soils of *F. microstachys*. Further investigation of plant biomass responses suggests that the observed shifts in coexistence outcomes arise because conditioned soils of *F. microstachys* substantially decrease conspecific growth (relative to unconditioned reference) in the immediate treatment, but this negative impact of conditioned soils is minimal after the time lag (Fig. 4B). Second, instead of representing the uncertainty of stabilization and fitness difference as orthogonal error bars, we visualized the distribution of random samples (i.e., using the same approach as in Yan *et al*., 2022 but without the summary pie chart). Our results show that visualizing the spread of predicted outcomes reveals informative patterns: for example, in the case of *A. wrangelianus* and *P. erecta*, the diagonal distribution in the delayed-with-litter treatment arises due to the larger variation in the impact imposed by *P. erecta* soils (Fig. 4). We echo recent calls for more careful considerations when calculating and visually presenting the uncertainty of predicted competitive outcomes (Terry & Armitage, 2023).

We incorporated litter dynamics in our experiment by preserving dead plant individuals in the pot during the temporal gap between the conditioning and response phases. This design corresponds well with our annual plant system, where litter input comes from natural plant senescence after the growing season. In other systems, different modes of plant death can generate litter dynamics that shape soil microbial communities differently. For instance, wind disturbances that uproot entire plants generate a substantial pulsed input of litter, potentially benefiting saprotrophic microbes but adversely affecting obligated root-associated microbes (Cowden & Peterson, 2013, Nagendra & Peterson, 2016). Conversely, herbivores and anthropogenic activities (e.g., clear-cutting) primarily remove aboveground parts while leaving belowground components intact in the soil. Such dynamics result in less aboveground litter, yet the remaining root tissues may continue to support arbuscular mycorrhizal fungi (Pepe *et al*., 2018). Recent studies have highlighted that aboveground and belowground litter differentially impact plant–soil microbe interactions. For example, Aldorfová *et al*. (2022) showed that root litter negatively affected plant performance due to enhanced pathogen transmission while shoot litter modified soil nutrient levels without significantly affecting plant growth. This complicated interplay between different belowground processes underscores the importance of including litter dynamics in microbially mediated plant– soil feedback studies (Ke *et al*., 2015, Veen *et al*., 2019).

We speculate that the impact of a temporal delay extends beyond that shown in our experiment. First, while we measured plant biomass performance following common plant-soil feedback studies, the temporal delay may modify microbial effects on other plant demographic rates. For example, seed survival and germination of Californian annuals take place in dry soil after the Mediterranean summer. Following recent calls for studying microbial impact beyond biomass, we encourage future research to study the impact of microbial persistence on the early-stage seed-to-seedling transition (Dudenhöffer *et al*., 2018, Miller *et al*., 2019, Krishnadas & Stump, 2021). Furthermore, how persistent are conditioned microbial effects after plant death is an important question for many systems, not limited to systems with strong seasonality and non-overlapping generations. For instance, in systems with sparse vegetation cover (e.g., foredunes), a conditioned patch may be left vacant for extended periods due to dispersal limitation. In more complex systems with vertical structures (e.g., forests), one may argue that soil microbes mostly impact seedling survival on conditioned soil beneath the canopy of a living adult. However, the persistence of microbial effects following adult death and canopy opening can influence the performance of seedlings, thereby determining which species can successfully reach the canopy. Therefore, investigating the persistence of conditioned microbial effects is likely more critical to community dynamics than previously recognized (Ke & Levine, 2021).

## Conclusion

In our effort to bridge the gap between greenhouse experiments and natural ecosystems, we demonstrate the feasibility and importance of adjusting experimental schedules to provide a more realistic representation of natural systems. In light of our findings, we propose that in annual plant systems with non-overlapping generations, the intricate interplay of natural seasonality and litter dynamics prevent the direct extrapolation of plant–soil microbe interactions from one growing season to the next. Our results reveal that the modification of plant–soil feedback following plant death is complex and varies between species pairs, thereby hindering generalizations based on studies that did not consider these factors. With the ongoing shifts in plant phenology and seasonal patterns due to climate change (Rudgers *et al*., 2020), predicting plant community dynamics requires the explicit incorporation of the temporal aspects and natural history elements into plant–soil feedback research (Ke *et al*., 2021).

## Acknowledgements

We acknowledge the Chumash peoples as the traditional land caretakers where the soils and plants for this project were collected. We recognize that Stanford University occupies the ancestral land of native peoples in the Bay Area. This work was performed (in part) at the University of California Natural Reserve System (Sedgwick Reserve, Reserve DOI: 10.21973/N3C08R), under Application #45441. We thank Kate McCurdy and the staff of Sedgwick Reserve, the Kraft lab at UCLA for providing seeds, and Jonathan Levine for his feedback on the experimental design. This research was supported by the California Native Grasslands Association. PJK is supported by the Taiwan MOE Yushan scholar program (MOE-110-YSFAG-0003-001-P1) and Taiwan NSTC (111-2621-B-002-001-MY3). MLW is supported by the USA NSF Graduate Research Fellowship (NSF GRFP DGE 1656518).

## Authorship Contribution

SXO, GSK, and PJK conceptualized and designed the study. SXO performed the experiment and sample processing. All authors contributed to data analyses and writing.

## Data Availability Statement

All primary data for microbial communities and the greenhouse experiment, as well as R scripts for data analysis, are available on GitHub (https://github.com/ousuzanne/PoundsofSoftFeta) for the review process. Microbial 16S rRNA and ITS sequences are available under the BioProject ID PRJNA1106794. All data and R scripts will further be archived on Zenodo upon acceptance.

## Supporting Information

The following Supporting Information is available for this article:

**Methods S1**. Supporting methods for soil microbial community characterization

**Fig. S1**. Principal coordinates analysis for the bacterial community composition

**Fig. S2**. Principal coordinates analysis for the fungal community composition

**Fig. S3**. Effects of soil microbial inocula on plant biomass in all response and control treatments

## Supporting Methods S1

### Synthetic spike-in

Using the synthetic spike-in method from Tkacz *et al*. (2018), we bought plasmids with p-Spike P for prokaryotic 16S (https://www.addgene.org/101172/) and p-Spike F for fungal communities (https://www.addgene.org/101174/). We plated the plasmids on Luria Broth (LB) media with carbenicillin for the ampicillin selection marker and incubated overnight in 25*^◦^*C. We then picked a colony and inoculated 10 mL of LB broth containing carbenicillin and incubated at 30*^◦^*C / 120rpm measuring CD on a nanodrop machine at 2 hour intervals until CD concentration = 1. Using zymoPURE Plasmid miniprep kit, we eluted plasmid DNA and measured DNA concentration using High Sensitivity dsDNA Qubit Assay (Thermofisher, Waltham, MA). We loaded a subsample of the eluted plasmid DNA for gel electrophoresis to check for the correct plasmid size.

### Amplicon sequencing

For bacterial metabarcoding, we amplified the highly variable (V4) region of the 16S rRNA gene using primers 515F (5*^1^*- TCG TCG GCA GCG TCA GAT GTG TAT AAG AGA CAG GTG YCA GCM GCC GCG GTAA -3*^1^*) and 806R (5*^1^*- GTC TCG TGG GCT CGG AGA TGT GTA TAA GAG ACA GGG ACT ACN VGG GTW TCT AAT -3*^1^*). For fungal metabarcoding, we amplified the fungal ITS1 region using primers based on the ITS1F (5*^1^*- AAT GAT ACG GCG ACC ACC GAG ATC TAC ACG GCT TGG TCA TTT AGA GGA AGT AA -3*^1^*) and ITS2 (5*^1^*- CAA GCA GAA GAC GGC ATA CGA GAT – [INDEX] – CGG CTG CGT TCT TCA TCG ATGC -3*^1^*), where [INDEX] is a sample-specific 12-nt error-correcting Golay barcode. Illumina adapters on each 5’ end of the primers were used to attach unique Nextera XT indexes for sample identification. First step PCR consisted of 3.2*µ*L of PCR-grade water, 5*µ*L of Meridian Bioscience MyTaq HS Red Mix (Bioline, Tunton, MA), 0.4*µ*L each of forward and reverse primers, and 1*µ*L of extracted DNA. PCR cycles were: 95*^◦^*C for 2 min, 35 cycles of 95*^◦^*C for 20 sec, 50*^◦^*C for 20 sec, 72*^◦^*C for 50 sec, and a final extension at 72*^◦^*C for 10 min with storage at 4*^◦^*C. We confirmed amplification by gel electrophoresis. Second step PCR consisted of 3.2*µ*L of PCR-grade water, 5*µ*L of Meridian Bioscience MyTaq HS Red Mix (Bioline, Tunton, MA), 0.4*µ*L each of Nextera XT index primers 1 and 2, and 1*µ*L of first step PCR product. We confirmed amplification by gel electrophoresis and purified amplicons using Sera-Mag Speedbeads (Sigma-Aldrich, St. Louis, MO). We quantified DNA concentration using High Sensitivity dsDNA Qubit Assay (Thermofisher, Waltham, MA) and pooled evenly across samples to a concentration of 4nM. The final DNA concentration was quantified using BioAnalyzer and sequenced on an Illumina MiSeq sequencer (2 X 300 cycle sequencing kit, Illumina, San Diego, CA) with a 15% PhiX spike-in at the Stanford Genomic Sequencing Service Center.

### Metabarcoding analysis

Reads were demultiplexed and assigned to samples using Illumina bcl2fastq conversion software. We processed ITS1 and 16S samples separately. We trimmed raw amplicon sequences using Cutadapt (Martin, 2011). We used the DADA2 pipeline (Callahan *et al*., 2016a) to merge paired-end sequences, quality filter, remove chimeric reads, and cluster sequences into amplicon sequence variants (ASVs). Potential contaminants were filtered using the decontam package (version 1.22.0; Davis *et al*., 2018), which removed 1 fungal ASV and 37 bacterial ASV. We used the SILVA database (Quast *et al*., 2012) for 16S taxonomic assignment and the UNITE database (Nilsson *et al*., 2019) for ITS taxonomic assignment. We removed any ASV that was present in *≤* 5 samples or whose relative abundance was *<* 0.01 across all samples. We also removed samples with extremely small or large read counts (i.e., more or less than 5x the average number of reads across all samples). We rarefied samples to 5000 sequencing reads.

## Supporting Figures

**Figure S1.**
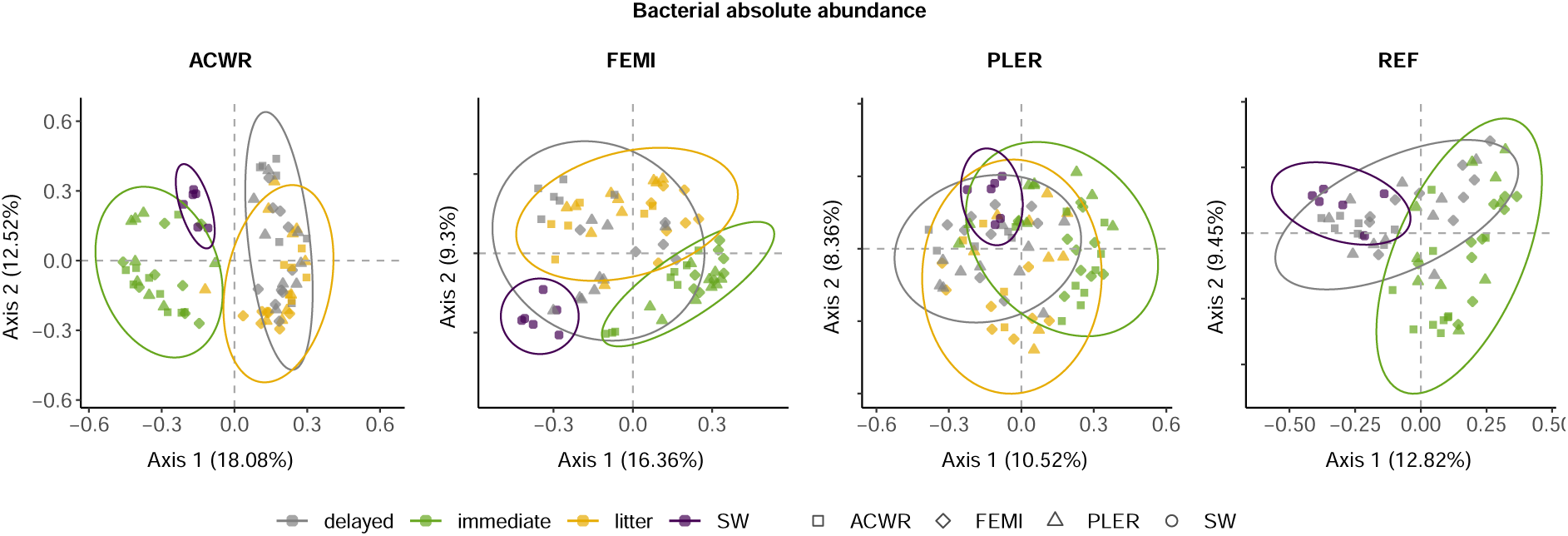
Principal coordinates analysis (PCoA) for the bacterial community composition sequenced at the end of the response phase. Each panel represents a different inoculum source (conditioning host plant). From left to right: *Acmispon wrangelianus* (ACWR); *Festuca microstachys* (FEMI); *Plantago erecta* (PLER); unconditioned Sedgwick Reserve field soil as reference soil (REF). Each point represents the microbial community sampled from a seedling at the end of the response phase and the shape represents its species identity. Colors represent the three response treatments: immediate (light green), delayed without litter (grey), and delayed with litter (brown). As the two delayed treatments shared the same reference soil controls, we omitted one of the delayed treatment in the rightmost panel. Purple circles (labeled as SW) represent soils collected from Sedgwick Reserve at the beginning of the experiment (i.e., without the growth of any conditioning or responding individual) and were added for visualization purposes.

**Figure S2.**
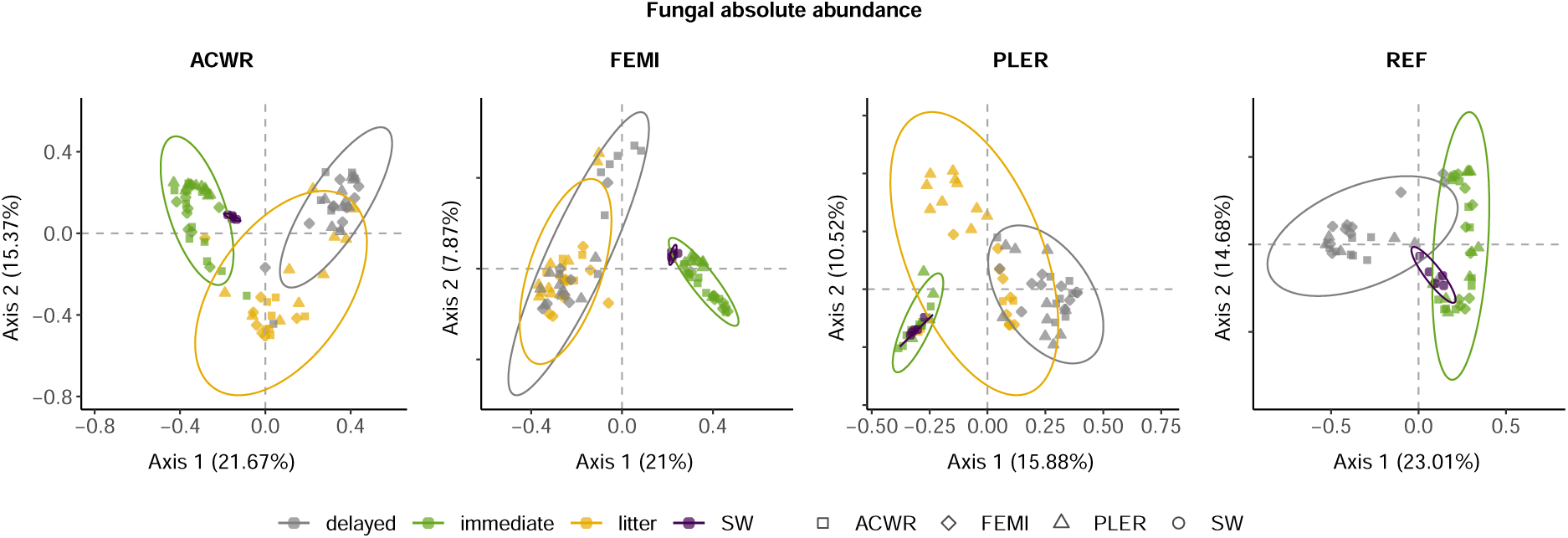
Principal coordinates analysis (PCoA) for the fungal community composition sequenced at the end of the response phase. Each panel represents a different inoculum source (conditioning host plant). From left to right: *Acmispon wrangelianus* (ACWR); *Festuca microstachys* (FEMI); *Plantago erecta* (PLER); unconditioned Sedgwick Reserve field soil as reference soil (REF). Each point represents the microbial community sampled from a seedling at the end of the response phase and the shape represents its species identity. Colors represent the three response treatments: immediate (light green), delayed without litter (grey), and delayed with litter (brown). As the two delayed treatments shared the same reference soil controls, we omitted one of the delayed treatment in the rightmost panel. Purple circles (labeled as SW) represent soils collected from Sedgwick Reserve at the beginning of the experiment (i.e., without the growth of any conditioning or responding individual) and were added for visualization purposes.

**Figure S3.**
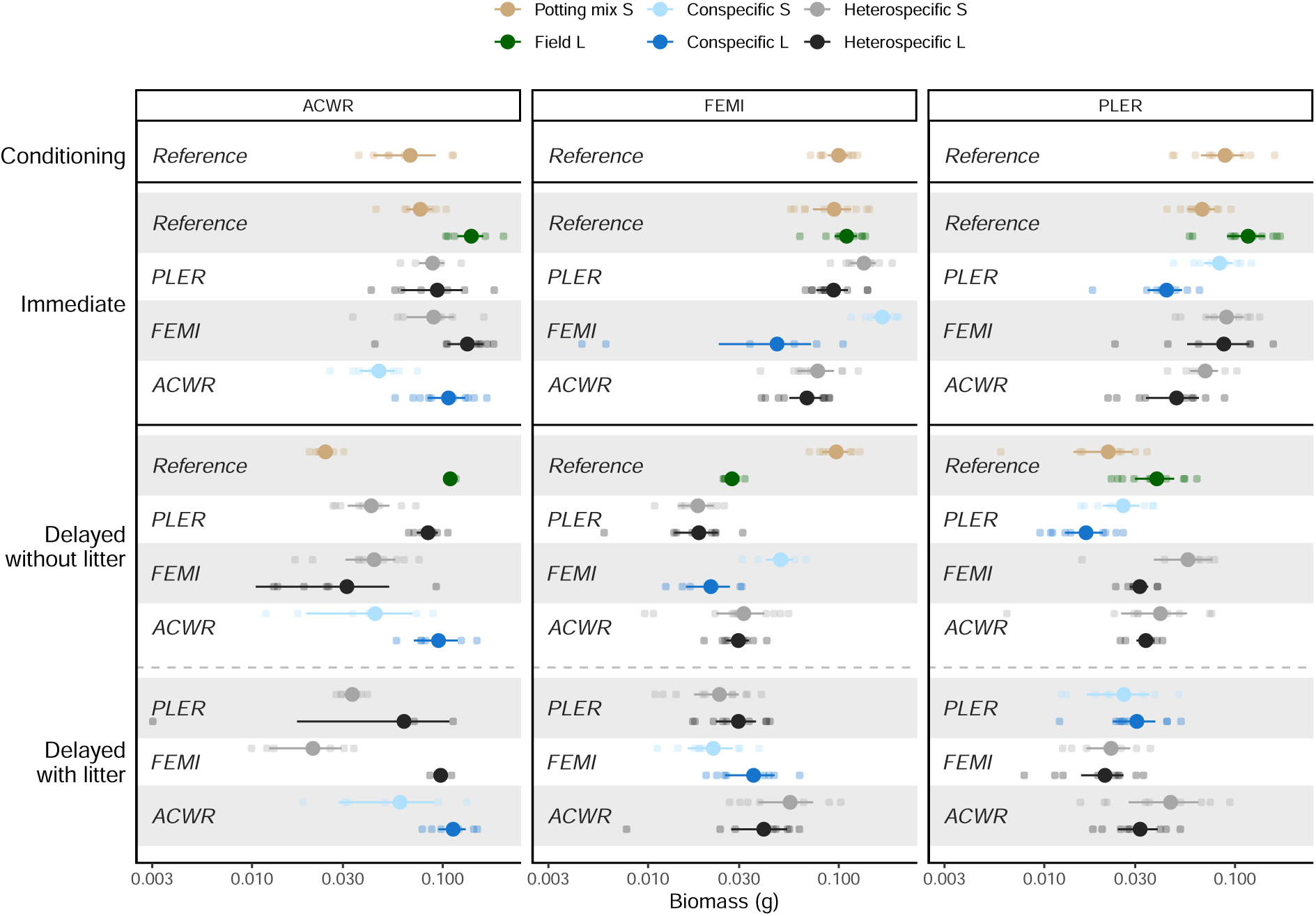
Effects of soil microbial inocula on plant biomass in all response and control treatments for (A) *Acmispon wrangelianus* (ACWR), (B) *Festuca microstachys* (FEMI), and (C) *Plantago erecta* (PLER). Capital “S” indicates sterilized soils and “L” indicates live un-sterilized soils. Colors represent different soil inocula: sterilized potting mix (brown), unconditioned field soil (green), soil conditioned by conspecifics (blue), soil conditioned by conspecifics but sterilized (light blue), soils conditioned by heterospecifics (dark grey), and soils conditioned by heterospecifics but sterilized (light grey). Note that the two delayed treatments shared the same field reference and sterilized potting mix controls. The three plant-conditioned soil inocula are ordered (from bottom to top) as follows: ACWR, FEMI, and PLER. Larger symbols indicate the mean biomass, error bars show 2 *×* SEM, and small points show each individual biomass.

